# An edge-centric model for harmonizing multi-relational network datasets

**DOI:** 10.1101/2021.01.07.425450

**Authors:** Joshua Faskowitz, Jacob C. Tanner, Bratislav Mišić, Richard F. Betzel

**Affiliations:** Department of Psychological and Brain Sciences, Indiana University, Bloomington, IN 47405; Program in Neuroscience, Indiana University, Bloomington, IN 47405; Cognitive Science Program, Indiana University, Bloomington, IN 47405; School of Informatics, Computing, and Engineering, Indiana University, Bloomington, IN 47405; McConnell Brain Imaging Centre, Montréal Neurological Institute, McGill University, Montréal, Quebec, Canada; Network Science Institute, Indiana University, Bloomington, IN 47405

## Abstract

Functional and structural connections vary across conditions, measurements, and time. However, how to resolve multi-relational measures of connectivity remains an open challenge. Here, we propose an extension of structural covariance and morphometric similarity methods to integrate multiple estimates of connectivity into a single edge-centric network representation. We highlight the utility of this method through two applications: an analysis of multi-task functional connectivity data and multi-measure structural networks. In these analyses, we use data-driven clustering techniques to identify collections of edges that covary across tasks and measures, revealing overlapping mesoscale architecture. We also link these features to node-level properties such as modularity and canonical descriptors of brain systems. We further demonstrate that, in the case of multi-task functional networks, edge-level features are consistent across individuals yet exhibit subject-specificity. We conclude by highlighting other instances where the edge-centric model may be useful.

## INTRODUCTION

The human brain is a network and can be modeled as a graph of nodes and edges, which represent neural elements and their pairwise functional interactions or structural links, respectively [1–3]. Using computational tools from graph theory and network science, the structure and function of brain networks can be probed, revealing key organizational and operational principles, including small-worldness [4, 5], modular structure [6–9], spatial constraints [10–12], and hubs and rich clubs [13–15].

Despite widespread application of network models within neuroscience, key challenges remain. In fact, even the most fundamental question of “what does it mean for two parts of the brain to be connected?” is not completely resolved [16–18]. In general, the answer to this question will vary with data modality, the time at which the measurement is made, the state in which the brain is situated, and the cognitive operation being performed. User preferences also play an important and somewhat arbitrary role – e.g. should functional connectivity be measured with full or partial correlations? – as do study aims – e.g. can connectivity be correlational or does it need to reflect directed, causal relationships [19]? This lack of consensus surrounding how connectivity is defined also presents issues with interpreting, contextualizing, and comparing results across studies. Although defining connections is central to every network neuroscience study, how to reconcile the multi-relational nature of brain connectivity is largely unaddressed.

One particularly useful strategy for measuring connectivity is based on interregional morphometric similarity (alternatively referred to as “anatomical covariance”) [20–23]. Calculating these measures involves first defining a set of features for each brain region and then, for pairs of regions, calculating their distance (or proximity) from one another in feature space. In most applications, feature vectors encode population-level variation of some morphological measurement, e.g. a region’s cortical thickness. The resulting similarity matrix is therefore representative of an entire cohort, making this approach poorly suited for studying individual differences. More recently, however, this approach was extended by defining feature vectors that represent a spectrum of structural and morphological measures for brain regions, e.g. their myelination status, curvature, and volume [24–27]. This extension makes it possible to define a similarity matrix at the level of individual subjects.

Both group- and subject-level morphometric networks are defined at the regional level, meaning that network nodes represent brain regions. Recently, we proposed an alternative, edge-centric method for constructing brain networks, emphasizing a network’s edges as its irreducible units and generating a higher-order network [28– 32]. Edge-level networks have a number of useful properties not shared by node-level networks, e.g. when clustered they naturally resolve overlapping clusters. More generally, edge-centric networks can be analyzed to reveal rich edge-level topology and present novel features that can potentially be used as biomarkers [33–35].

Here, we combine the edge-centric framework with an extension of morphometric similarity to integrate multirelational connectivity data into one edge-level network model. Although this approach is flexible and can be applied to any set of connectivity measurements, we focus on two specific examples that highlight its utility for neuroscience. First, we combine rest and task-evoked functional connectivity to study multi-task reconfiguration and second, we integrate multiple measurements of structural connection weights. In these examples, we demonstrate that edge covariance networks can be decomposed into clusters to reveal groups of edges that cohesively modulate their weights across tasks or vary similarly across different weighting schemes of structural networks.

## RESULTS

Throughout this manuscript we analyze neuroimaging data from the Human Connectome Project [36] and a dense phenotyping study of a single individual [37]. In the first section, we analyze functional neuroimaging data from both datasetes. In the second section, we analyze structural networks estimated using HCP data. Details of the acquisition, processing, and network construction procedures can be found in **Materials and Methods**.

### Edge covariance estimation

The presence/absence and weight of connections can be estimated using many different methods. To consolidate this variability into a single model, we propose a novel edge-centric analog to morphometric similiarity networks. In this section, we briefly summarize the procedure (see **Materials and Methods** for a complete description).

Traditionally, interregional morphometric similarity networks are constructed by estimating the pairwise similarity between brain regions’ structural and morphological features [20, 24]. This approach can be extended to the level of edges by estimating the connection weight between pairs of nodes based on different modalities, e.g. structural or functional connectivity, using different measurements, e.g. full/partial/regularized correlation, spectral coherence, mutual information, etc., or under different conditions, e.g. task or rest. These multi-relational estimates of a connection’s weight can be assembled into a feature vector (Fig. 1a) and the similarity of this vector calculated with respect to any other edge in the network (Fig. 1b). Repeating this procedure for all *m* edges yields an edge-by-feature matrix (Fig. 1c) that can be further transformed into an edge covariance (or edge similarity) matrix by computing the similarity between all pairs of edge feature vectors (Fig. 1d). We note that the edge covariance matrix is only modestly correlated with the edge functional connectivity matrix described in our previous reports (*r* = 0.48; *p <* 10^−15^; Fig. S1a) and its topology not clearly driven by spatial relationships (correlation of edge covariance with surface areas of quadrilateral traced out by the four nodes involved in an edge pair, *r* = −0.02; Fig. S2b).

**FIG. 1.**
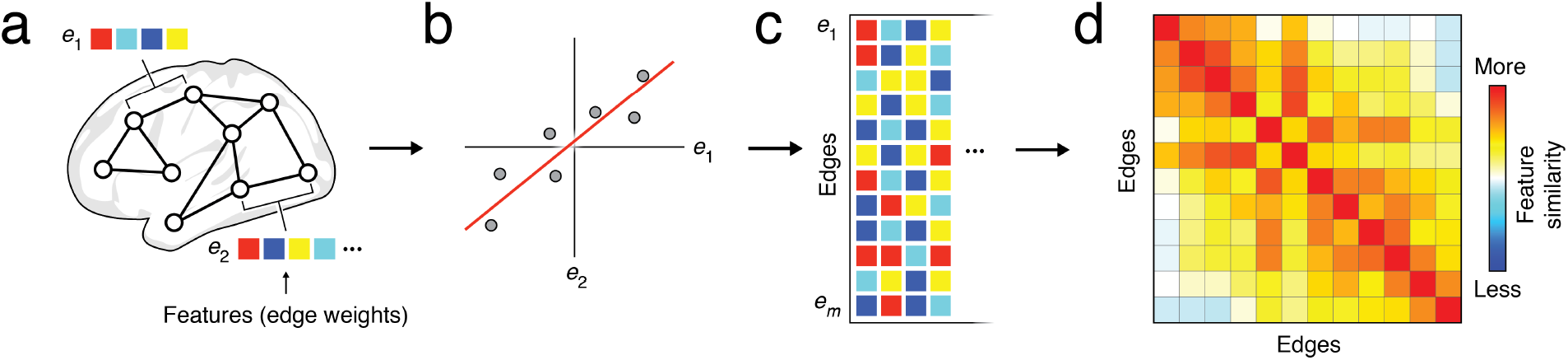
Edge covariance network estimation. (*a*) Each edge is associated with a set of features that represent estimates of its weight using different modalities and measures and/or conditions. (*b*) We can calculate the connection weight between any two edges, e.g. as the similarity of their feature vectors. In this diagram, points represent features, e.g. different means of weighting an edge. (*c*) Scaled to the entire brain, this procedure results in a matrix of edge feature vectors. (*d*) An edge covariance network can then be estimated as the matrix of all pairwise similarity values.

Edge covariance matrices differ from morphometric networks in several key ways. First, edge covariance matrices have dimensionality of ℝ^*m*×*m*^, whereas morphometric networks have dimensionality of ℝ^*n*×*n*^, where *n* and *m* are the number of nodes and edges in the network, respectively. Whereas morphometric networks compare measurements taken at the physical nodes, edge covarience networks compare information about the interrelationships between nodes. Note, also, that edges’ feature vectors are flexible and can be defined multiple ways. For instance, they could represent connectivity estimates made using different imaging modalities, while subjects complete different tasks, or different measurements of connection weight made on the same network dataset. They could even reflect connectivity estimates at different points in time, in which case edge covariance is nearly identical to edge functional connectivity, which we analyzed in previous studies [28–30].

### Task-based edge covariance matrices

It is well-documented that, when estimated using Pearson correlation, functional connection weights are subtly yet systematically shifted when subjects perform cognitively demanding tasks, driving the brain into increasingly integrated states [38–42]. However, the principles by which edge weights reconfigure remain unclear [43]. For instance, are there certain classes of connections whose weights are similar across different tasks? Are some connections selectively modulated in a small number of tasks while others fluidly reconfigure across many? Here, we address these questions by constructing an edge covariance matrix from group-representative resting-state and task-evoked functional connectivity. Specifically, we define a set of features for each edge based on its weight during the seven tasks included in the HCP dataset (EMOTION, GAMBLING, LANGUAGE, MOTOR, RELATIONAL, SOCIAL, working memory (WM)) as well as rest (see **Materials and Methods** for details on preprocessing).

#### Modular structure

Are there clusters of edges that reconfigure similarly across multiple cognitive domains? In our first analyses we address this question by clustering the multi-task edge covariance matrix. To do this, we first calculated functional connectivity as the full matrix of correlation coefficients for the 92 subjects that pass quality assurance for all resting-state and the seven task scans (Fig. 2a). For each edge, we extracted its correlation weight for each task, averaged these values over all subjects, and standardized (z-score) their values to generate a feature vector for each edge (Fig. 2b). In principle, we could then estimate the edge covariance network by calculating the pairwise similarity of edges’ feature vectors (Fig. 2c; we show examples of these correlations in Fig. S1). The resulting edge-by-edge matrix could be treated as a network and clustered using any of the familiar community detection methods [44]. However, for computational ease, we opted to directly cluster the matrix of feature vectors. Although, there exist many methods for clustering data, we were again motivated practically and used k-means with the correlation distance metric. We varied the number of clusters from *k* = 2 to *k* = 25, repeating the algorithm 250 times at each parameter setting. We selected the optimal number of clusters to be the *k* at which the mean similarity of estimated clusters was both large and exhibited low variance. These heuristics revealed both *k* = 2 and *k* = 6 as good candidates. We focused on *k* = 6 for all subsequent analyses and derived consensus clustered from the 250 estimates (Fig. 2b-d; see **Materials and Methods** for details).

**FIG. 2.**
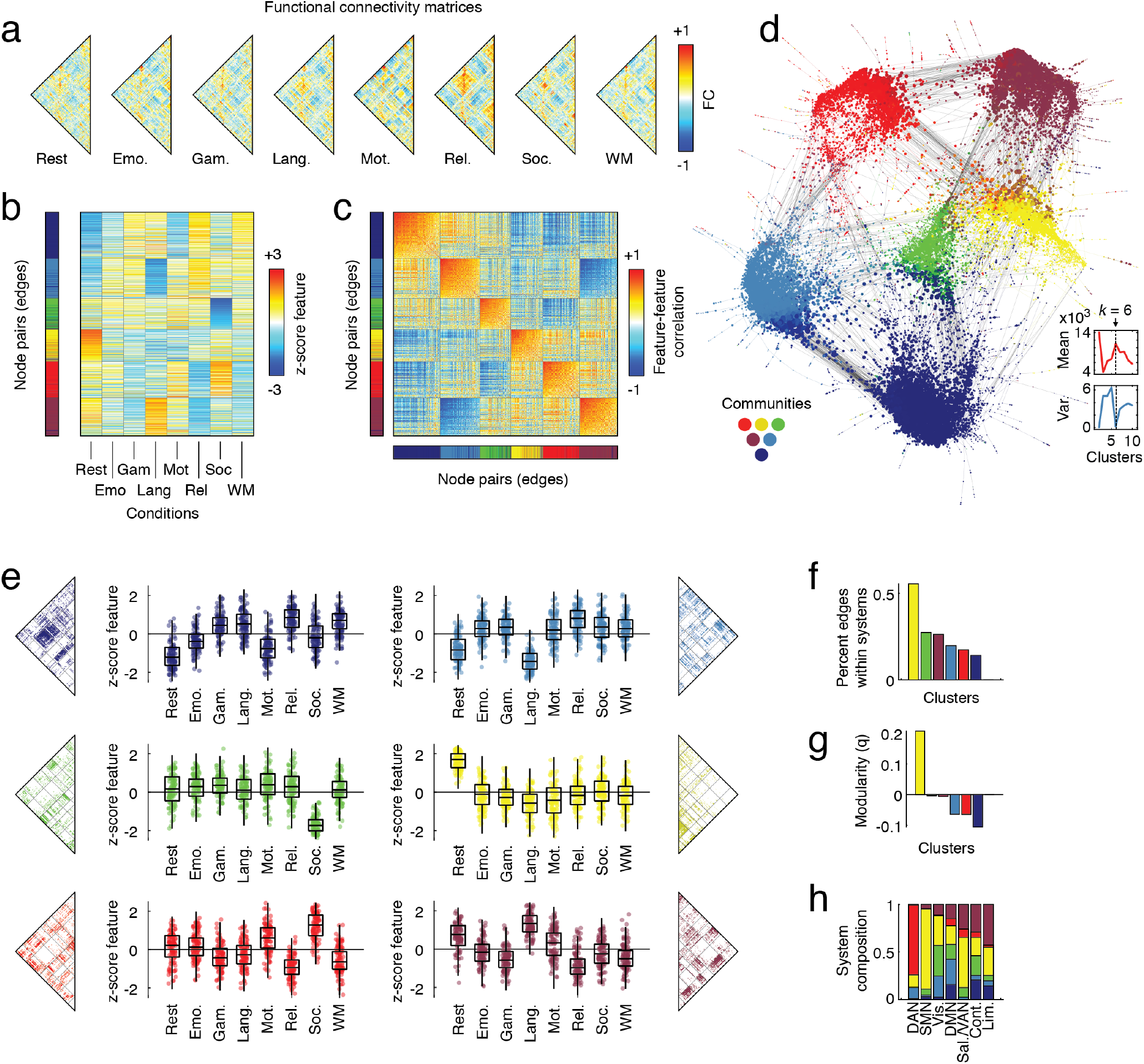
Edge covariance matrices for multi-task functional connectivity data. (*a*) Functional connectivity networks estimated during eight conditions. (*b*) Normalized edge feature matrix with rows sorted by edge clusters. (*c*) Edge covariance matrix sorted by edge clusters. (*d*) Force-directed layout of edge covariance matrix (union of minimum spanning tree and strongest 0.01% connections. Inset shows mean and variance of z-score Rand index as a function of the number of clusters, *k*, with peaks and troughs at *k* = 6. (*e*) “Connectional fingerprints” for each of the six clusters depicting normalized weights of edges across different task conditions. (*f*) For each cluster, the fraction of its connections that fall within canonical brain systems. (*g*) The modularity, *q*, of each cluster. (*h*) Composition of within-system edges in terms of edge cluster assignments.

From the clustering results, we defined a connectional fingerprint for each of the six clusters (Fig. 2e). Our results allow us to make several interesting observations. First, we find that two clusters are largely associated with specific tasks. Clusters three (green) and four (yellow) include connections that tend to decrease and increase their weights during the SOCIAL task and at REST, respectively, but only subtly modulate their connections during all other conditions (Fig. 2e). Interestingly, cluster four included many within-system connections, indicating that the cohesive functional connectivity that supports the emergence of brain systems is largely rest-specific. Other clusters exhibit more complicated fingerprints that involve modulation of edge weights across multiple tasks. Cluster one (dark blue), for instance, includes connections between the default mode and somatomotor network that decrease their weights during REST and MOTOR tasks, decrease their weights modestly during EMOTION and SOCIAL tasks, and increase weights during all others.

The clustering results can also be viewed as node-level networks, themselves. That is, the set of edges assigned to each cluster can be treated as node-by-node network and can be analyzed separately. We show the upper triangle of each such network alongside their respective fingerprints in Fig. 2e. We also calculate and report a select set of characteristics for each network, including the extent to which their connections fall within brain systems, their modularity, and their system composition, i.e. the canonical brain systems that are concentrated within each cluster [45]. We find that cluster four (yellow) is the only cluster that has greater than 50% of its connections concentrated within traditionally defined brain systems, whereas all other clusters are dominated by between-system connections. Not surprisingly, this same cluster was the only one that exhibited a positive modularity (Fig. 2g). Lastly, we considered the composition of traditionally defined systems in terms of their within-system edges’ cluster assignments (Fig. 2h). We found that dorsal attention and somatomotor networks were dominated by clusters five and four, respectively, and that other systems contained mixtures of many different clusters. Notably, only default mode and control networks included edges associated with all six clusters.

Note that for the analysis reported here, we calculated standardized edge features based on functional connectivity estimates using all frames, irrespective of their in-scanner motion. To confirm that our analyses are not obviously biased by motion, we repeated this procedure but based on functional connectivity where we dropped high-motion frames (relative motion *>* 0.1 mm). We found that the group-representative standardized feature vectors were highly correlated (*r* = 0.998), suggesting that motion is not an obvious confound in these analyses. A second potential issue concerns the relative duration of the scans and the number of samples used to estimate functional connectivity. To circumvent any possible issues, we repeated our analyses after recalculating functional connectivity using the same number of frames in all tasks (equal to the minimum number of frames in any task). Again, we found that the feature vectors were highly correlated (*r* = 0.979), suggesting that the number of samples is not an obvious confound for these analyses.

Collectively, these results suggest that edge covariance patterns can be used to reveal clusters of edges that reconfigure across tasks. These results hint that task-evoked changes in functional connectivity are fundamentally low-dimensional [46] and can be characterized by a small number of template patterns or fingerprints [47].

#### Overlap

In the previous section, we extracted and clustered edge-level features from resting-state and task-evoked functional connectivity, resulting in a non-overlapping partition of edges into clusters. However, edge clusters induce overlapping clusters from the perspective of individual nodes. Here, we characterize these overlap patterns to identify brain regions and systems that participate in few or many edge clusters.

To extract measures of edge cluster overlap, we first reshape cluster labels into the upper triangle of a node-by-node matrix (Fig. 3a). Then, for each node, we can calculate the fraction of its edges that are affiliated with each of the six clusters (Fig. 3b). These projections of edge-level clusters onto nodes yield an overlapping view of clusters. That is, rather than forcing nodes to have all-or-nothing affiliation with clusters, i.e. non-overlapping clusters, each node can be affiliated fractionally with multiple clusters through its edges’ cluster assignments. Interestingly, we find that the affiliation measures exhibit non-random topography and align closely with canonical systems. For instance, cluster four (yellow) largely involves edges with stub nodes falling within somatomotor and visual networks (Fig. 3b). Similarly, cluster one, two, and six involve edges with stubs in default mode, control, and visual systems, respectively (Fig. 3b). Interestingly, clusters three and five exhibit similar patterns of fractional participation (*r* = 0.50, *p* = 8.2 × 10^−14^), with both involving edges linked to visual, dorsal attention, and control networks. Although the projections of these clusters onto brain regions are similar, the clusters involve disjoint sets of edges.

**FIG. 3.**
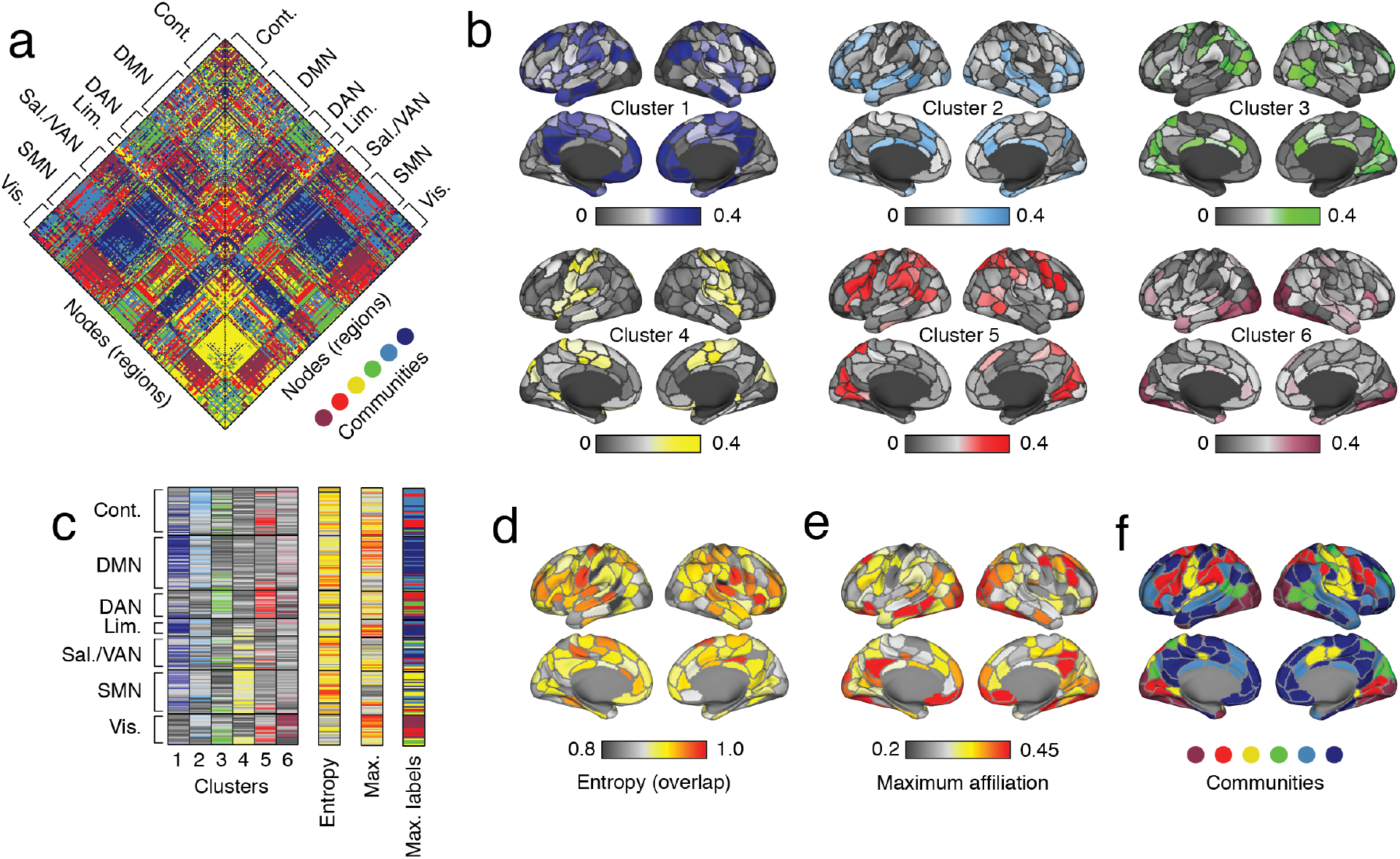
Edge cluster projections and overlap. (*a*) Node-by-node matrix with edges labeled according to the cluster to which they were assigned. (*b*) Projections of edge clusters onto brain regions. (*c*) Matrix representation of projections (left) and the overlap level, the max affiliation, and the label associated with the max affiliation of each node. (*d*) Topographic representation of overlap patterns. (*e*) Maximum affiliation of each brain region to any of the clusters. (*f*) Label of the cluster to which each brain region was maximally affiliated.

We can also compute several summary statistics based on the affiliation measure (see **Materials and Methods**). First, as in [29, 48], we calculated the normalized entropy of each nodes’ edge cluster assignment distribution. Intuitively, if a node’s edges are assigned to a single cluster (or a small number of clusters), this entropy measure is close to 0. However, if a node’s edges are assigned to many clusters and evenly distributed, then the entropy is close to 1. Additionally, we also calculated each node’s maximum affiliation, i.e. the cluster to which the greatest number of its edges were assigned (Fig. 3c). Interestingly, we found high levels of entropy (overlap) across the cortex, with values approaching 1 in many regions and with 98.5% of regions (197/200) exhibiting an entropy greater than 0.8 (Fig. 3d). At the system level, no system exhibited significantly different levels of entropy (spin test). In fact, when we ranked nodes according to the entropy, we found that high-entropy nodes (top 25%) could be found in every system except for visual.

Next, we calculated the maximal affiliation of each region to *any* of the six clusters. We found no evidence that the maximum affiliation was significantly elevated or depressed in any system and that, as in the previous section, regions ranked in the top 25% based on their maximum affiliation could be found in every brain system. However, when we examined the clusters to which regions maintained their maximum affiliation, we found that the resulting partition was highly similar to the canonical brain systems reported in [45] (adjusted Rand index, *ARI* = 0.75; spin test *p <* 10^−3^), suggesting that system-level architecture may play a role in determining task-evoked patterns of functional network reconfiguration [49, 50].

#### Repeatability

Finally, we assessed the reliability and repeatability of normalized edge features, the primary ingredient for calculating edge covariance matrices. This procedure involved several analyses carried out using functional imaging data from two datasets: the Human Connectome Project and dense phenotyping data from a single individual that included thirty scan sessions with both rest and task conditions of similar duration [37].

Using HCP data, we first tested whether normalized feature matrices were similar across individuals by computing the inter-subject similarity of observed matrices with those obtained after permuting subjects’ functional connectivity matrices using a spatially-constrained test [51] (Fig. 4a). Note that while the same permutations were applied to all scans of the same individual, different permutations were used for different individuals. In general, we found that subjects were more similar to one another than expected by chance (Fig. 4b; t-test; *p <* 10^−3^).

**FIG. 4.**
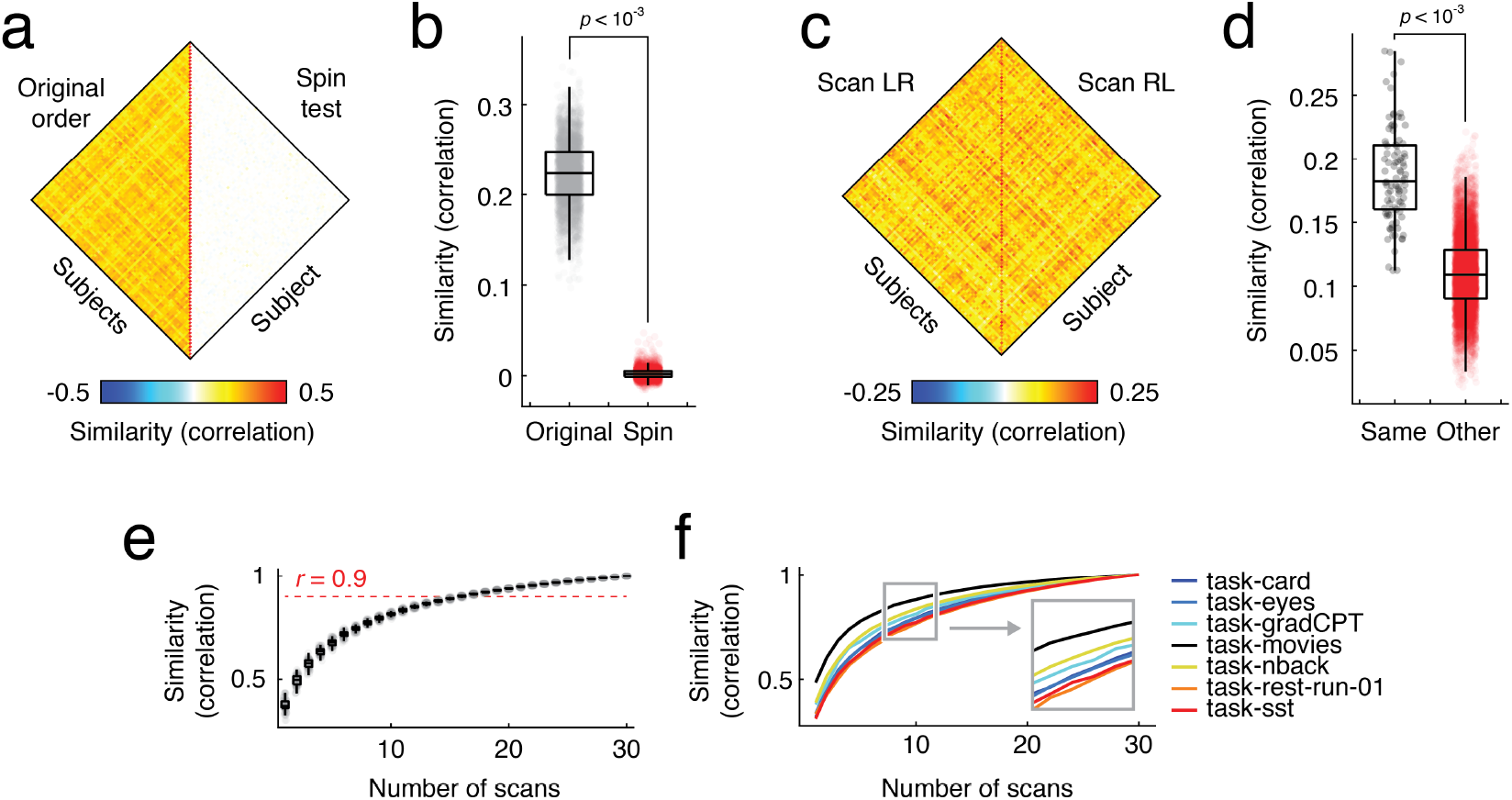
Subject specificity and repeatability of normalized edge feature matrices. (*a*) Within-subject similarity of normalized edge feature matrices compared with spin test (nodal permutation that preserves spatial relationships among brain regions) *b*) Boxplot showing distributions of similarity values for original and spin test data. (*c*) Similarity of normalized edge feature matrices separately estimated using left-to-right and right-to-left phase encoding data. (*d*) Distribution of within-subject and between subject similarity. (*e*) Similarity of subsampled normalized edge feature matrices to grand average. (*f*) Similarity of specific tasks.

Next, we tested whether normalized feature matrices were subject-specific. To do this, we generated separate feature matrices using the left-to-right and right-to-left (LR and RL, respectively) phase encoding scans. We then computed the similarity of these matrices to one another (Fig. 4c). If features were subject-specific, we would expect to find that elevated similarity between the LR and RL matrix from the same subject compared to matrices from different subjects. In the similarity matrix, this effect would yield a bright red trace along the diagonal. We found that this was the case, suggesting that normalized feature matrices are not only similar across the HCP cohort, but are also unique to each subject (Fig. 4d; t-test; *p <* 10^−3^).

Finally, we wanted to know how much data was necessary to obtain an accurate estimate of normalized edge features. To do this, we analyzed data collected from a single individual over 30 scan sessions during which that individual underwent resting-state scans along with multiple task conditions (card-sorting, n-back task, gradual-onset continuous performance task, stop-signal, card-guessing, reading the mind in the eyes, movie-watching) [37]. With multiple observations/scan sessions from the same individual performing an identical set of tasks, this dataset is useful for assessing the effect that different amounts of data have on the veracity of the feature matrix.

To this end, we calculated functional connectivity matrices for each scan and condition (Pearson correlation as the measure of connectivity) and subsequently computed a grand average across all scans. As before, we vectorized each edge’s weight across all conditions, standardized the weights of each edge, and arranged the standard scores into a matrix. Then, we repeated this procedure using random samples of scan sessions, first with each scan session on its own, then with pairs of sessions, then triplets, quartets, and so on. Each time, we used more data to obtain an average, gradually building towards using the full amount of data available across all scan sessions. For each random sample, we also calculated the similarity of its feature matrix to the grand average matrix (Fig. 4e). We found that the similarity to the grand average increased monotonically as more samples were used to estimate feature matrices, yielding diminishing returns. To achieve a level of similarity (Pearson correlation) of *r* = 0.9, data from 16 scan sessions was required (482 × 16 = 7712 samples). We also calculated the similarity of each column in the feature matrix separately and found that the movie scan stabilized more quickly than the others.

In summary, our findings suggest that standardized edge feature matrices are similar across subjects, exhibit high-levels of subject specificity, and small increases in the amount of data leads to rapid increases in the stability of their estimates, followed by a protracted regime of diminished returns. These observations suggest that feature matrices and edge covariance matrices may be useful substrates for investigating multi-modal differences in connectivity patterns across individuals [52, 53].

### EDGE COVARIANCE MATRICES FOR STRUCTURAL CONNECTIVITY

In the previous section, we introduced a novel edge-centric analog of morphometric similarity, prioritizing an analysis of functional MRI data acquired during restingstate and task conditions. However, the edge covariance matrix framework is general and can be applied to other imaging modalities. Here, we demonstrate this flexibility by applying this technique to structural connectivity networks reconstructed from diffusion MRI using tractography (see **Materials and Methods** for more details). One of the key challenges in studying structural brain networks is that there exists multiple methods for weighting network edges [54, 55]. Some weights are based on tractography measure, e.g. streamline counts, normalized number of streamlines, etc.. Others are based on microstructural properties of the reconstructed tracts, e.g. the mean fractional anisotropy along a streamline’s path, its mean diffusivity, diffusion kurtosis, etc..

Here, we use the edge-covariance method to reconcile disparate measurements of structural connectivity, focusing on: streamline counts, normalized counts, number of streamlines, fractional anistropy, mean diffusivity, and diffusion kurtosis (see **Materials and Methods** for detailed definitions), and combining them into a single edge-centric model. However, this procedure requires some additional preprocessing steps. First, unlike functional connectivity, structural brain networks are fundamentally sparse. This means that most edges do not exist and, accordingly, we have no measure of their connection weight. In this analysis, we omit those edges. Second, structural connectivity edge weights vary from one another by orders of magnitude, making them especially difficult to compare. For instance, streamline counts are integer values and can vary from a value of 0 to a maximum determined by the number of seeds, with typical values as large as 10^3^ or 10^4^. In contrast, measures like fractional anisotropy are bounded to the interval [0, 1]. To account for these disparities, we opted to resample edge weights within modalities from a zero-mean and unit variance Gaussian distribution while preserving weight ranks, a procedure used previously to ensure normally distributed edge weights [56]. All analyses reported in the following sections were carried out using a group-representative structural network [57].

The group-representative structural connectivity matrix was sparse (*δ* = 0.30; *m* = 6036 undirected edges) (Fig. 5a, left; ordered by nodal clusters; see Fig. S3 for topographic representation of edge clusters). Of the existing edges, we extracted the Gaussian resampled weights, combined them into a normalized feature matrix, and clustered this matrix using k-means. We used the same heuristic as in the previous section to select the number of clusters and found that repeated runs of the algorithm converged to a highly similar solution at *k* = 5. We analyzed the consensus partition obtained from these data here (edges with cluster labels are shown in Fig. 5a, right).

**FIG. 5.**
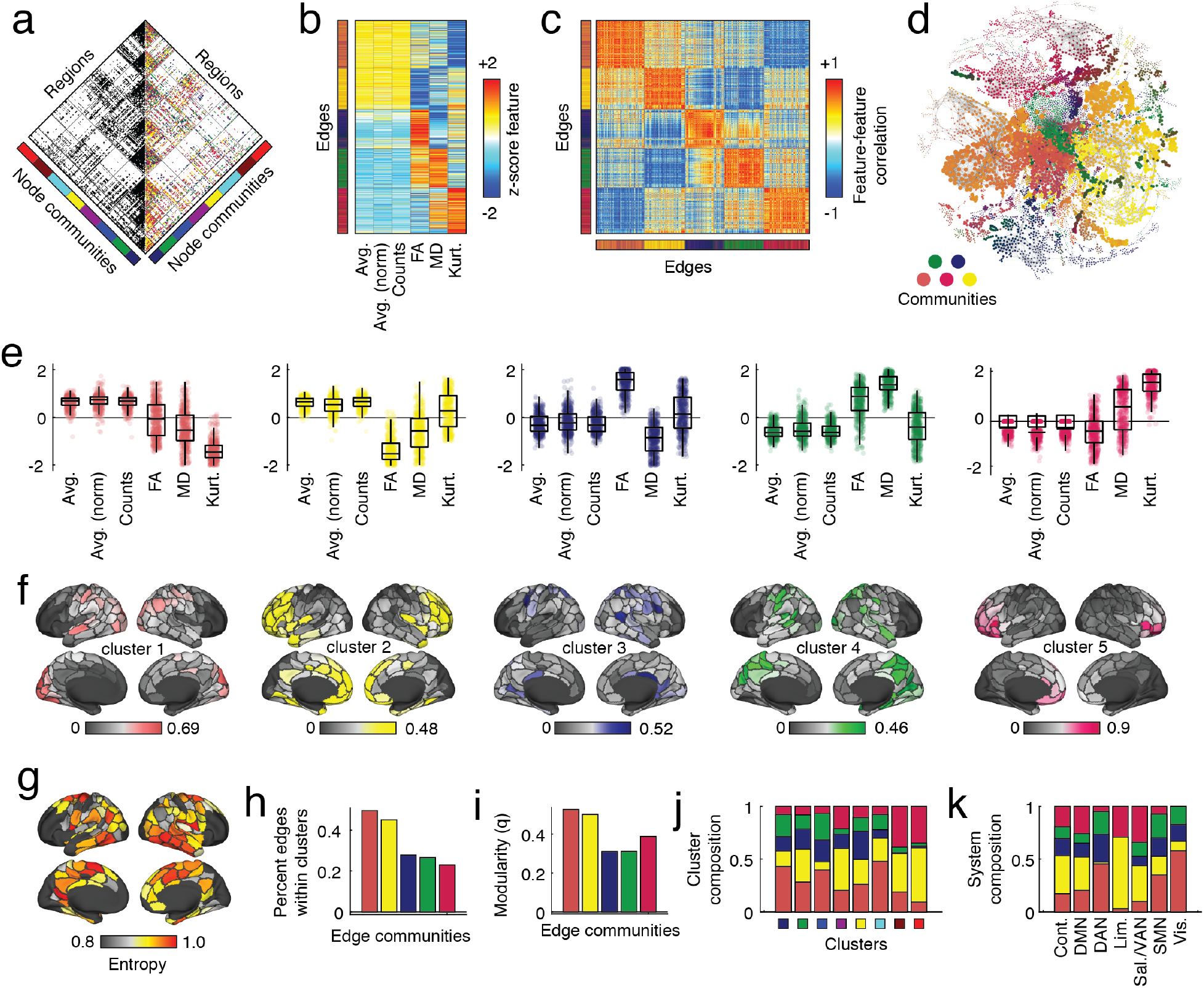
Edge covariance matrices for structural connectivity data. (*a*) Binary structural connectivity mask (left) and edge cluster assignments (right). (*b*) Normalized edge feature vector with rows sorted by edge clusters. (*c*) Edge covariance matrix sorted by edge clusters. (*d*) Force-directed layout of edge covariance matrix (minimum spanning tree plus 0.001% strongest connections). (*e*) Connectional fingerprints. (*f*) Projections of edge clusters onto brain regions. (*g*) Edge cluster entropy (overlap). (*h*) Fraction of connections associated with each edge cluster that fall within node clusters. (*i*) Modularity of each edge cluster. (*j*) Edge cluster composition of node clusters. (*k*) Edge cluster composition of canonical brain systems.

The edge clusters divided the normalized feature vectors into five clusters (Fig. 5b) and resulted in an edge covariance matrix with high levels of within-cluster similarity (Fig. 5c,d). The first two clusters were comprised of edges that exhibited elevated streamline-based connectivity (average, normalized average, and total streamline count) and reductions in edge weights of diffusion kurtosis (cluster one) and fractional anisotropy (cluster two) (Fig. 5e). In contrast, clusters three, four, and five exhibited modest decreases in streamline-based connectivity measures, but elevated connection weights for fractional anisotropy (cluster three), mean diffusivity (cluster four) and kurtosis (cluster five) (Fig. 5e). Note also that, broadly, edge clusters partition structural links into groups of dissimilar lengths (Euclidean distance; Fig. S4).

Interestingly, when we project edge clusters onto the cortical surface, we find distinct topographic representations aligned broadly with canonical brain systems (Fig. 5f). Cluster one overlapped closely with visual and dorsal attention networks, cluster two with control and default mode, cluster three with somatomotor and visual networks, cluster four with somatomotor, default mode, and dorsal attention, and cluster five with control and salience/ventral attention network. The edge clusters also induce typical patterns of overlap (normalized entropy; Fig. 5g). Similar to the entropy measured using functional edge covariance clusters, the overlap is reduced in visual cortex but is elevated to some extent in all other systems.

Finally, we analyzed the edge clusters patterns as subgraphs. We found that different edge clusters one and two contained edges that tended to fall within nodal clusters (Fig. 5h) and exhibited elevated modularity (Fig. 5i). Edge clusters also were also distributed non-randomly within node clusters (Fig. 5j) and canonical brain systems (Fig. 5k).

Collectively, these results suggest that edge covariance matrices can be used to integrate multiple connectivity modalities into a single edge-level network. Our findings further suggest that edge weights covary with one another in sometimes complicated patterns, but that by combining multiple measures into the same model, we can better understand these relationships and their links to network topology and the brain’s system-level organization.

## DISCUSSION

In this manuscript, we introduce a method for integrating multi-relational datasets into a unified edge-level network representation. We apply this framework in two contexts: first to study task-evoked changes in functional connectivity and second to combine different weighting schemes for structural connections. Our results suggest that edge-level models capture repeatable and subject-specific patterns of connectional covariance and identify non-overlapping clusters of edges that fluctuate similarly across tasks and weights (analyses one and two, respectively). We also demonstrate that edge clusters can be projected to the level of brain regions, yielding distinct patterns of cluster overlap. Our work capitalizes upon and extends other recent edge-level models, further highlighting their viability for neuroscience.

### Edge covariance networks for neuroscience

Brain networks are made up of nodes and edges. Although there are many ways to divide a brain into nodes, it is generally agreed upon that each node should represent a distinct area or parcel of gray matter (although occasionally nodes are defined as sensors/electrodes). However, the question of how to define edges is more complicated, and represents a moving target that varies with imaging modality, condition, and/or time. Finally, the decision is somewhat arbitrary and usually left to the user’s discretion, leading to a multitude of possible weighting schemes.

Here, we present an edge-centric strategy for addressing this challenge. Rather than force a user to weight their network using only one of many possible sets of weights, we construct an edge-by-edge network whose organization reflects edges’ covariances with one another. This approach not only helps sidestep one of the key challenges in network neuroscience, but it offers many advantages, including the ability to categorize edges into sub-networks and to detect pervasively overlapping clusters.

To date, applications of edge-centric models in network neuroscience are uncommon and, in some cases, not wholly appreciated as being edge-centric. For instance, several early papers used sliding windows to estimate time-varying functional networks and track the correlation of edges with one another across time, inadvertently creating edge-level networks [58–60]. Other recent papers have deliberately adopted edge-centric approaches, usually citing [33, 34] as inspiration. In addition to the edge time series and edge functional connectivity models discussed earlier [28–32], others have generated edge-edge networks by embedding edges in a some metric space and using inter-edge distances to quantify the connection weight between edges [35].

Although these approaches all adopt an edge-centric perspective, their mathematical underpinnings differ as does the intuition they provide about brain network organization. Moving forward, an important avenue for future work involves systematically comparing these different approaches to better understand their relative advantages and disadvantages.

### Relationships with existing methods

The edge-centric model presented here is, to our knowledge, novel. However, it is similar to a number of existing methods. For instance, the general framework of computing covariances (or more accurately, correlations) between a set of feature vectors and treating the resulting matrix as a network has been studied at length in the form of structural covariance matrices [20], which are a special case where the features correspond to a regional measure of brain structure (usually cortical thickness) estimated across a population. This approach has been extended recently [24] by calculate a set of features for a given region, making it possible to estimate subject-specific structural covariance matrices (morphometric similarity matrices). Note, also that previous studies have investigated the covariance structure of other node-level properties using network measures. These include gene and transcriptomic similarity networks [61–63], microstructural profiles [64], co-atrophy patterns [65], meta-analytic co-activation networks [66], and time series features [67].

In our work we generate a similar construct but at the level of connections rather than regions. Specifically, we estimate features at the level of edges and compute the pairwise similarity between feature vectors for edge dyads, resulting in an edge-by-edge correlation matrix. This extension of connectomics from nodes to edges is in line with several recent papers [28, 35], which build on earlier work promoting edge-centric analysis of complex networks [33, 34]. In particular, our work here generalizes the methods from Faskowitz et al [28]. In that paper, the authors compute a measure called “edge functional connectivity” as the similarity of edge time series – measurements of the co-fluctuation between pairs of brain regions. In the context of the current paper, we can think of co-fluctuation time series as a set of edge-level features, and edge functional connectivity as a specific instance of an edge covariance matrix.

### Future directions

Our work opens up a multitude of opportunities for future work. Any dataset in which there exists multiple measurements of an edge’s weight can be immediately transformed into its corresponding edge covariance matrix. For instance, building on our analysis of multi-task edge covariance networks, one can apply this approach to data from individual subjects to investigate inter-individual differences in edge-level organization [68]. Note that while our approach requires multitask data in order to construct an edge covariance matrix, it would be straightforward to calculate subject-specific edge covariance matrices using data from only a single condition, e.g. the resting state, by computing multiple measures of connectivity and assessing edges covariances across these measures. Our work also helps resolve methodological issues associated with preprocessing pipelines. For instance, it is well known that decisions in preprocessing can yield dissimilar estimates of functional connectivity downstream in the pipeline [69, 70]. Rather than trying to identify the singularly optimal pipeline, our work could allow users to take the agnostic approach and treat functional connectivity estimates from *many* pipelines as features and compute an edge covariance network based on these estimates. For instance, one could include models that include and omit controversial steps like global signal regression [71, 72]. Yet another possible extension involves the construction of edge covariance matrices from multi-subject data. Using this approach one could examine patterns of inter-subject variability.

A final possibility is to consider multi-modal edge-covariance networks. For instance, one could include features for node pairs based on their structural and functional connectivity, matrix measurements derived from those matrices (e.g. topolgoical distance, communicability, search information, etc. [73]), spatial relationships (e.g. Euclidean distance or geodesic distance on cortical surface), the similarity of regions’ transcriptomic profiles [62, 74], or even morphometric similarity. Of course, care needs to be taken that edge weights are of comparable scale and that missing connections are dealt with appropriately. This type of model could help provide new insight into structure-function relationships [75] or, more generally, help us understand the statistical relationship between different connection modalities.

### Limitations

Here, we present an edge-centric model for resolving variable estimates of connection weights. The model operates by treating multi-relational connectivity data as a set of edge-level features and the edge-by-edge similarity matrix as the connectivity matrix for a network. However, in the effort to maintain a sense of agnosticism concerning edge weights, we nonetheless are forced to make a decision about weights in the edge covariance network. Here, we select the Pearson correlation to maintain consistency with the most common measure of functional connectivity, but note that other measures could be used.

A second related issue concerns the calculation of similarity between edges using only on a small number of features. It is understood that the sampling distribution around correlation coefficients narrows as the number of samples increases. Consequently, estimating edge covariance from a limited number of samples may hinder the ability to accurately calculate edge covariance matrices. In the future, several steps could be taken to mitigate this issue. For instance, the number of samples used to estimate similarity could be increased dramatically by including data from many subjects, rather than group-level data as was done here. Doing so requires also modeling the effect of individual differences, e.g. some subjects may exhibit increases or decreases in their baseline connectivity so that when edge weights are standardized, all data from these subjects appears elevated or depressed relative to the other subjects. Nonetheless, this strategy would help narrow the sampling error associated with the edge similarity measures and improve statistical power for identifying “true” edge-level interactions.

A final limitation concerns the analysis of structural connectivity. Here we calculate edge covariance based on features of sparse structural edges (physical white-matter tracts between brain regions), resulting in an edge covariance matrix whose dimensionality does not match that of the functional connectivity covariance matrix. In principal, however, we could match their dimensions by transforming the sparse structural connectivity matrix into a fully-weighted matrix by calculating relational measures between pairs of brain regions, e.g. the similarity of their connectivity profiles, the length and cost of the shortest topological path between them, or their capacity to communicate with one another [73]. Here, however, we focus on the structural connections alone.

## MATERIALS AND METHODS

### Datasets

The Human Connectome Project (HCP) dataset [36] consisted of resting state and task functional magnetic imaging (fMRI) data, as well as diffusion magnetic resonance imaging data (dMRI) from 100 unrelated adult subjects (54% female, mean age = 29.11 ± 3.67, age range = 22-36). These subjects were selected as they comprised the “100 Unrelated Subjects” (U100) released by the Human Connectome Project. After excluding subjects based on data completeness and quality control (see **Quality Control**), the final fMRI subset utilized included 92 subjects (55% female, mean age = 29.32 ± 3.66, age range = 22-36). The final dMRI subset utilized included 95 subjects (56% female, mean age = 29.29 ± 3.66, age range = 22-36). The study was approved by the Washington University Institutional Review Board and informed consent was obtained from all subjects. A comprehensive description of the imaging parameters and image prepocessing can be found in [76, 77]. Images were collected on a 3T Siemens Connectome Skyra with a 32-channel head coil. Subjects underwent two T1-weighted structural scans, which were averaged for each subject (TR = 2400 ms, TE = 2.14 ms, flip angle = 8°, 0.7 mm isotropic voxel resolution). Subjects underwent four resting state fMRI scans, and 14 task fMRI scans (2 of each task; EMOTION, GAMBLING, LANGUAGE, MOTOR, RELATIONAL, SOCIAL, working memory (WM)) over a two-day span. The fMRI data was acquired with a gradient-echo planar imaging sequence (TR = 720 ms, TE = 33.1 ms, flip angle = 52°, 2 mm isotropic voxel resolution, multiband factor = 8). Each resting state run duration was 14:33 min, with eyes open and instructions to fixate on a cross. Task scans ranged from 2:16 to 5:01 min in duration, with 5 to 10 blocks per task. Details about each task can be found in [78]. Finally, subjects underwent two diffusion MRI scans, which were acquired with a spin-echo planar imaging sequence (TR = 5520 ms, TE = 89.5 ms, flip angle = 78°, 1.25 mm isotropic voxel resolution, b-vales = 1000, 2000, 3000 s/mm^2^, 90 diffusion weighed volumes for each shell, 18 b = 0 volumes). These two scans were taken with opposite phase encoding directions and averaged.

The single subject dataset included resting state and task fRMI collected over a 10-month period at Yale University. The data was originally collected as part of a study investigating the influence of tasks on functional parcellation estimation [37]. The subject (male, age = 56 at onset of study) underwent 33 scanning sessions. The subject provided written informed consent in accordance with a protocol approved by the Human Research Protection Program of Yale University. Images were collected on two identically configured Siemens 3T Prisma scanners with a 64-channel head coil. In each session, the subject underwent two resting state fMRI scans and six task scans (N-back, gradual-onset continuous performance, stop-signal, card guessing, reading the Mind in the eyes, movies). The fMRI data was acquired with a gradient-echo planar imaging sequence (TR = 1000 ms, TE = 30 ms, flip angle = 55°, 2 mm isotropic voxel resolution, multiband factor = 5). Each resting state run duration was 6:49 min, and task scans were approximately 6 min in duration. A T1-weighted structural scan (MPRAGE) was also acquired for the subject during the first session (TR = 2400 ms, TE = 1.22 ms, flip angle = 8°, 1 mm isotropic voxel resolution).

### Image preprocessing

#### HCP functional preprocessing

Functional images in the HCP dataset were minimally preprocessed according to the description provided in [76]. Briefly, these data were corrected for gradient distortion, susceptibility distortion, and motion, and then aligned to a corresponding T1-weighted (T1w) image with one spline interpolation step. This volume was further corrected for intensity bias and normalized to a mean of 10000. This volume was then projected to the *32k fs LR* mesh, excluding outliers, and aligned to a common space using a multi-modal surface registration [79]. The resultant cifti file for each HCP subject used in this study followed the file naming pattern: `* REST{1,2} {LR,RL} Atlas MSMAll.dtseries.nii`.

#### Single subject functional preprocessing

Functional images for the single subject dataset were preprocessed using *fMRIPrep* 1.3.2 [80], which is based on Nipype 1.1.9 [81]. The following description of *fMRIPrep*’s preprocessing is based on boilerplate distributed with the software covered by a “no rights reserved” (`CC0`) license. Internal operations of *fMRIPrep* use Nilearn 0.5.0 [82], ANTs 2.2.0, FreeSurfer 6.0.1, FSL 5.0.9, and AFNI v16.2.07. For more details about the pipeline, see the section corresponding to workflows in *fMRIPrep*’s documentation.

The T1-weighted (T1w) image was corrected for intensity non-uniformity with `N4BiasFieldCorrection` [83, 84], distributed with ANTs, and used as T1w-reference throughout the workflow. The T1w-reference was then skull-stripped with a Nipype implementation of the `antsBrainExtraction.sh` workflow, using NKI as the target template. Brain surfaces were reconstructed using `recon-all` [85], and the brain mask estimated previously was refined with a custom variation of the method to reconcile ANTs-derived and FreeSurfer-derived segmentations of the cortical gray-matter using Mindboggle [86]. Spatial normalization to the *ICBM 152 Nonlinear Asymmetrical template version 2009c* [87] was performed through nonlinear registration with `antsRegistration`, using brain-extracted versions of both T1w volume and template. Brain tissue segmentation of cerebrospinal fluid (CSF), white-matter (WM) and gray-matter (GM) was performed on the brain-extracted T1w using FSL’s `fast` [88].

Functional data was slice time corrected using AFNI’s `3dTshift` and motion corrected using FSL’s `mcflirt` [89]. *wang2017evaluation* distortion correction was performed by co-registering the functional image to the same-subject T1w image with intensity inverted [90] constrained with an average fieldmap template [91], implemented with `antsRegistration`. This was followed by co-registration to the corresponding T1w using boundary-based registration [92] with 9 degrees of freedom. Motion correcting transformations, field distortion correcting warp, BOLD-to-T1w transformation and T1w-to-template (MNI) warp were concatenated and applied in a single step using `antsApplyTransforms` using Lanczos interpolation. Several confounding timeseries were calculated based on this preprocessed BOLD: framewise displacement (FD), DVARS and three regionwise global signals. FD and DVARS are calculated for each functional run, both using their implementations in Nipype [93]. The three global signals are extracted within the CSF, the WM, and the whole-brain masks. The resultant nifti file utilized for this study followed the file naming pattern `* space-T1w desc-preproc bold.nii.gz`.

#### HCP structural preprocessing

Diffusion images in the HCP dataset were minimally preprocessed according to the description provided in [76]. Briefly, these data were normalized to the mean b0 image, corrected for EPI, eddy current, and gradient nonlinearity distortions, corrected for motion, and aligned to the subject anatomical space using a boundary-based registration [92]. In addition to this minimal preprocessing, images were corrected for intensity non-uniformity with `N4BiasFieldCorrection` [83]. FSL’s `dtifit` was used to obtain scalar maps of fractional anisotropy, mean diffusivity, and mean kurtosis. The Dipy toolbox (version 1.1) [94] was used to fit a multi-shell multi-tissue constrained spherical deconvolution [95] to the diffusion data with a spherical harmonics order of 8, using tissue maps estimated with FSL’s `fast` [88].

Tractography was performed using Dipy’s `Local Tracking` module. Multiple instances of probabilistic tractography were run per subject [96], varying the step size and maximum turning angle of the algorithm. Tractography was run at step sizes of 0.25, 0.4, 0.5, 0.6, and 0.75 with the maximum turning angle set to 20°. Additionally, tractography was run at maximum turning angles of 10°, 16°, 24°, and 30° with the step size set to 0.5. For each instance of tractography, streamlines were randomly seeded three times within each voxel of a white matter mask, retained if longer than 10 mm and with valid endpoints, following Dipy’s implementation of anatomically constrained tractography [97], and errant streamlines were filtered based on the cluster confidence index [98].

#### Image quality control

For the HCP fMRI data, scans were filtered out based on motion criteria recommended elsewhere [69]. Where a spike is defined as relative root mean squared (RMS) movement of 0.25 mm or greater, scans were excluded if one of the following criteria was met: 1) greater than 15% of frames are spikes, 2) average RMS movement greater than 0.2 mm, or 3) any spike larger than 5 mm. Furthermore, data was visually inspected for artifacts. After applying these criteria, eight subjects lacked a full set of fMRI data. For the HCP dMRI data, scans were filtered based on the motion estimated by FSL’s `eddy` and motion estimated during the subject’s associated resting scans. Scans were filtered out if measurements for resting state fMRI and dMRI mean and median absolute deviation (MAD) of the RMS motion exceeded 1.5 times the interquartile range (in the adverse direction) of the measurement distribution. After applying these criteria, four dMRI datasets were excluded. Furthermore, data or one subject failed to complete the spherical deconvolution step. For the single subject data, scans were visually inspected using the fMRIPrep output. All fMRI data was retained. However, data of three fMRI runs could not be downloaded from the source data repository, due to technical difficulties.

### Functional and structural networks preprocessing

#### Parcellation preprocessing

The Schaefer parcellation [45] was used to delineate 200 regions on the cortical surface. For HCP fMRI data, the cifti version of this atlas in’32k fs LR’ space was used. For the HCP dMRI data and the single subject data, a volumetric parcellation was rendered in anatomical space using the spherical warp computed by FreeSurfer’s `recon-all` [85]. This warp is based on individual curvature and sulcal patterns.

#### Functional network preprocessing

Each preprocessed BOLD image (from both the HCP and single subject datasets) was linearly detrended, band-pass filtered (0.008-0.08 Hz), confound regressed and standardized using Nilearn’s `signal.clean` function, which removes confounds orthogonally to the temporal filters. The confound regression strategy included six motion estimates, mean signal from a white matter, cerebrospinal fluid, and whole brain mask, derivatives of these previous nine regressors, and squares of these 18 terms. Spike regressors were not applied. This 36 parameter strategy has been show to be a relatively effective option to reduce motion-related artifacts, particularly in tandem with the filtering of high motion subjects [69]. Following these preprocessing operations, the mean signal was taken at each node.

#### Structural network preprocessing

The number of streamlines between nodes of the volumetric parcellations was recorded for each tractography instance. Fractional anisotrophy, mean diffusivity, and mean kurtosis maps were sampled from the middle 80% of each streamline’s path, which were averaged within streamline and then across all streamlines between each pair of nodes. Streamline counts were normalized by dividing the count between nodes by the geometric average volume of the nodes. Since tractography was run nine times per subject, edge values had to be collapsed across runs. To do this, the weighted mean was taken, with weights based on the proportion of total streamlines at that edge. This operation biases edge weights towards larger values, which reflect tractography instances better parameterized to estimate the geometry of each connection.

### Edge covariance networks

In this subsection we describe the procedure for constructing a generic edge covariance matrix. We also provide additional details about how edge covariance matrices were constructed for the multi-task functional connectivity data and multi-measure structural connectivity data.

Brain networks are usually modeled such that there exist *n* nodes linked to one another by a set of *m* edges. We let **W** ∈ ℝ^*n*×*n*^ be a weighted connectivity matrix where *W_ij_* is the weight of the connection between nodes *i* and *j*. In general, there may be *n_W_* ways of weighting that connection. We denote this ensemble of weights as 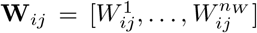 In practice, we z-score the elements of this vector, which we treat as a list of features for the edge *e_ij_*. These features can be assembled in matrix form as 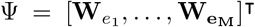 where *e_m_* is the feature vector for edge *m* 1*,…, m*. We can then calculate the matrix of feature correlations, Ω ∈ ℝ^*m*×*m*^, which we refer to as the edge covariance matrix, as 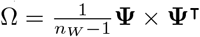.

### Clustering algorithm

We can also cluster the matrix of features, Ψ. Here, we use the traditional k-means algorithm with the correlation distance. We initialize the algorithm 250 times with different centroids, yielding slightly dissimilar solutions each time. To resolve the variability of solutions we use a modified consensus clustering algorithm. Briefly, this algorithm transforms each of the 250 estimates of clusters into a set of indicator vectors of size *M × k*, where *k* is the number of clusters. The elements of each indicator vector are one for points assigned to that cluster and zero otherwise. Then, we concatenate indicator vectors into a single matrix and perform an eigendecomposition, retaining the top *k* components which we then iteratively recluster using k-means with the same number of clusters. The algorithm stops when all 250 repeats arrive at the same solution.

Here, we use k-means to cluster edge feature vectors. We note that this choice is motivated practically and that, in general, any clustering algorithm could be applied to the feature matrix or the edge covariance matrix to obtain clusters. The output of the clustering algorithm is a partition of edges into clusters, such that 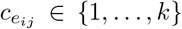 denotes the cluster to which edge *e_ij_* is assigned. We analyze these clusters in two formats. First, as the vector *C* ∈ ℝ^*m*×1^, but also as in matrix form as *D* ∈ ℝ^*n*×*n*^, with elements 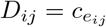.

#### Nodal affiliations

For each brain region, we can calculate its affiliation to any of the *k* clusters. To do this, we analyze the matrix representation of edge clusters. The affiliation of region *i* to edge cluster *c* ∈ {1*,…, k*} is defined as:

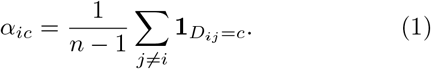

From this equation, we can also calculate the maximum affiliation of each brain region as 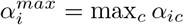.

This equation is modified for the sparse structural connectivity data. Rather than normalize by *n* – 1 (the total number of edges incident upon a node in a functional connectivity matrix), we normalize by each node’s degree, so that the affiliation reads:

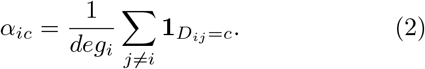

#### Entropy measures

We can also use the affiliation measurement to measure the extent to which a brain region participates in multiple clusters, i.e. its level of overlap. One possibility is to simply count the number of non-zero elements in each region’s affiliation vector, *α_i_* = [*α_i1_*,*…,α_ic_*]. This measure, however, could be biased by small values, e.g. instances where a node is affiliated primarily with a single cluster but maintains weak affiliations to the remaining clusters *via* single edges. Accordingly, we measure overlap using an entropy measure:

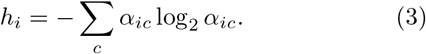

We then normalize this measure as 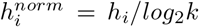, so that it is bounded to the interval [0, 1]. Intuitively, values close to 0 indicate regions whose edges maintain a consistent affiliation with a small number of clusters, while values close to 1 indicate regions whose edges are uniformly distributed over multiple clusters.

### Network measures

In the main text, we treat the set of edges assigned to a given cluster as a network. Like any other network, we can calculate properties of this network, including its cluster structure and modularity [99]. The modularity heuristic, *q*, measures the cohesiveness – within-cluster density of connections – compared to what would be expected under a chance model and is calculated as:

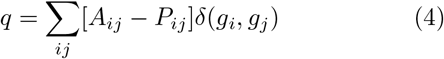

where *A_ij_* and *P_ij_* are the observed and expected weight of the edge *e_ij_*, *δ*(*x, y*) is the Kronecker delta function and is equal to 1 when *x* = *y* and 0 otherwise, and *g_i_* is the cluster assignment of node *i*. Here, we use the Louvain algorithm [100] to detect clusters in the edge networks and to quantify their modularity. For simplicity, with use a degree-preserving null model, so that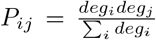. We also normalize *q* so that it is bounded to the interval [−1, 1] as follows: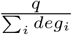.

#### Partition similarity

In general, there are many heuristics available for choosing the optimal number of clusters in a dataset. Here, we select the *k* at which the k-means algorithm consistently converges to a similar solution (low variance) and where the average partition similar is greater than nearby values of *k*. To do this, we compute for every *k* the similarity matrix comparing all pairs of the 250 detected partitions. As a measure of similarity, we use the z-score Rand index [101]:

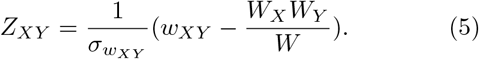

In this equation, *W* is the total number of node pairs in the network; *W_X_* and *W_Y_* are the total number of pairs within clusters in partitions *X* and *Y*; *w_XY_* is the number of pairs assigned to the same module in both *X* and *Y*; and *σ_XY_* is the standard deviation of *w_XY_*. The value of *Z_XY_* reflects the similarity between *X* and *Y* after accounting for chance.

Here, we compute the similarity between all pairs of partitions detected at each *k* and calculate the mean and variance. We select the optimal *k* to be the value at which the mean achieves a local maximum and where the variance is low, i.e. where the algorithm repeatedly converges to a similar solution.

#### Structural connectivity clusters

To estimate nodal clusters from structural connectivity, we used an extension of modularity maximization that accounts for spatial relationships, which are known to exert a strong pressure on structural connections [12]. Briefly, this approach involves reframing the modularity equation so that *P_ij_* depends on the distance between nodes *i* and *j*. To do this, we run a rewiring model 1000 times to estimate the value of *P_ij_* for all edges. This model exactly preserves each node’s degree and, to an arbitrary level of precision, preserves the network’s edge length distribution, and hence its total cost of wiring [102]. The resulting clusters can be interpreted as groups of nodes that are more densely connected to one another given their degree and the brain’s wiring cost constraints.

## DATA AVAILABILITY

All imaging data come from publicly-available, open-access repositories. Human connectome project data can be accessed *via* https://db.humanconnectome.org/app/template/Login.vm. after signing a data use agreement. The single subject data can be accessed *via* OpenNeuro at http://doi.org/10.18112/openneuro.ds002372.v1.0.0.

## ACKNOWLEDGEMENTS

This material is based upon work supported by the National Science Foundation under Grant No. 076059-00003C (RFB).

---

**FIG. S1.**
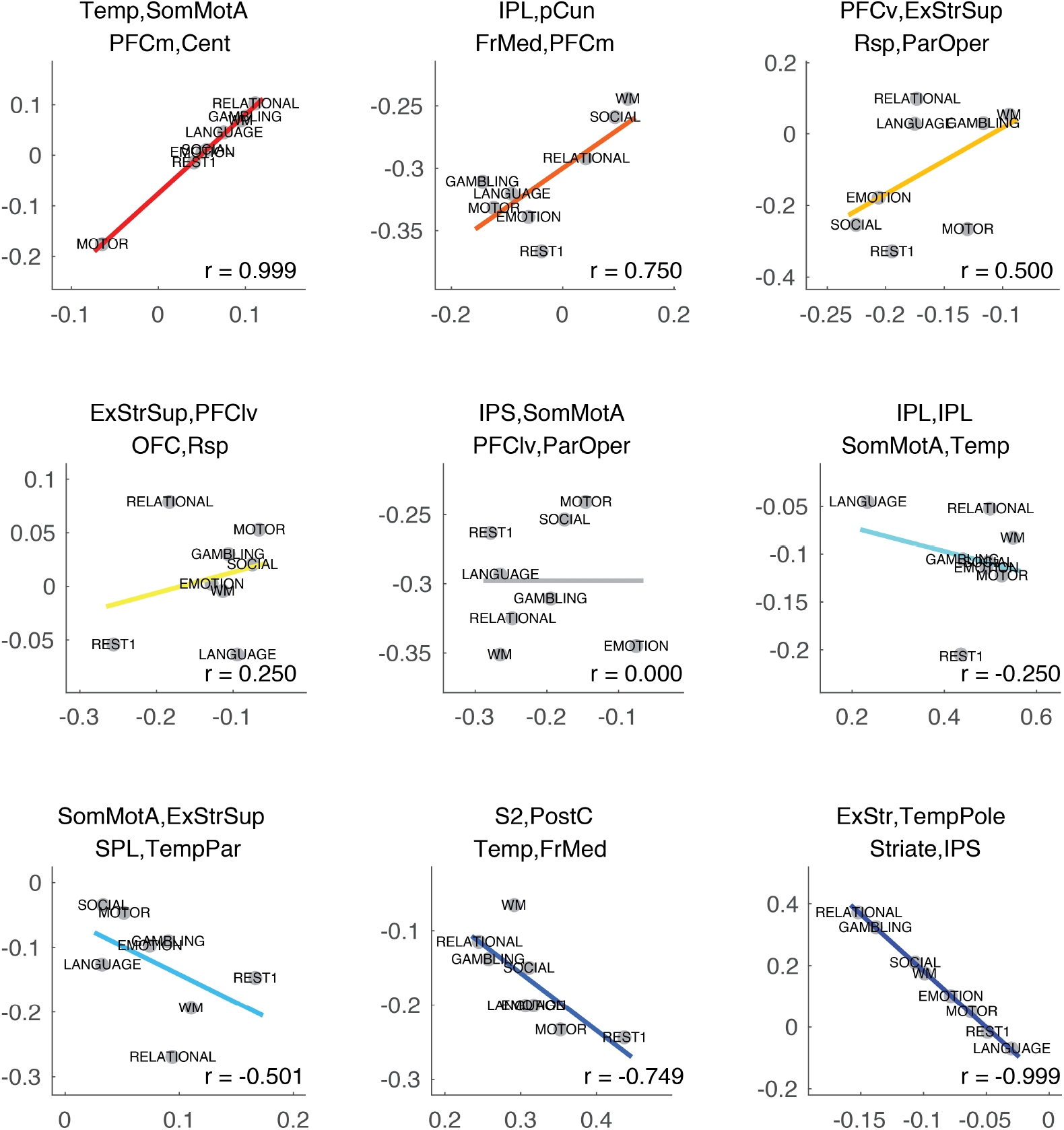
Examples of feature correlations for multi-task functional connectivity dataset. We show example scatterplots of normalized edge weights between pairs of edges. The stub regions (endpoint) of each edge are displayed at the top of each plot.

**FIG. S2.**
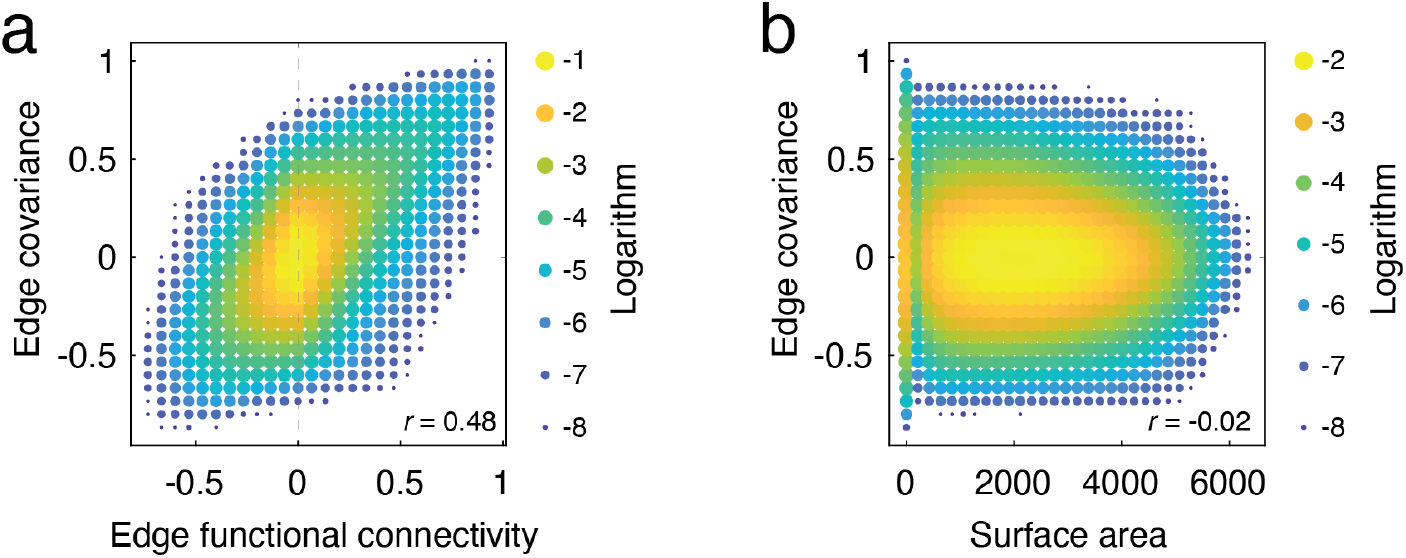
Relationship of edge covariance with edge functional connectivity and surface area. (*a*) Two-dimensional histogram of edge covariance weights and edge functional connectivity as estimated in [28]. (*b*) Two-dimensional histogram of edge covariance weights and the surface area of the quadrilateral formed by the four nodes comprising the two edge pairs.

**FIG. S3.**
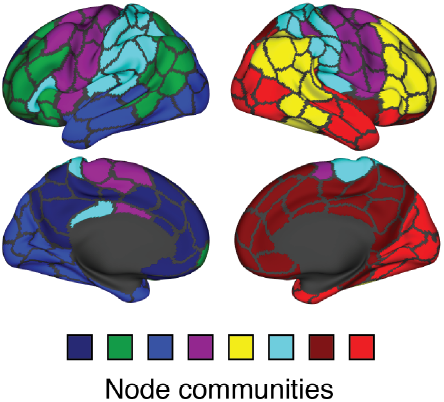
Topographic distribution of communities estimated from structural connectivity data.

**FIG. S4.**
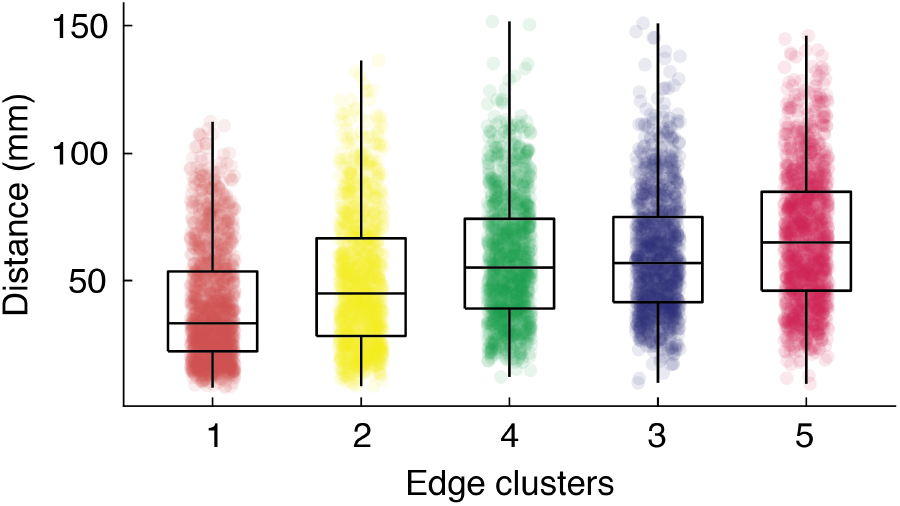
Length profiles of structural edge clusters. Boxplot of edge lengths (Euclidean distances) of edges assigned to the structural edge clusters. All clusters are significantly different from one another except for cluster three with four and three with five (*p <* 6.6 × 10^−10^; FDR corrected with false discovery rate fixed at *q* = 0.05.)

